# Network analysis of chromophore binding site in LOV domain

**DOI:** 10.1101/2022.12.10.519884

**Authors:** Rishab Panda, Pritam Kumar Panda, Janarthanan Krishnamoorthy, Rajiv K. Kar

## Abstract

Photoreceptor proteins are versatile toolbox for developing biosensors for optogenetic applications. These molecular tools get activated upon illumination of blue light, which in turn offers a non-invasive method for gaining high spatiotemporal resolution and precise control of cellular signal transduction. The Light-Oxygen-Voltage (LOV) domain family of proteins is a well-recognized system for constructing optogenetic devices. Translation of these proteins into efficient cellular sensors is possible by tuning their photochemistry lifetime. However, the bottleneck is the need for more understanding of the relationship between the protein environment and photocycle kinetics. Significantly, the effect of the local environment also modulates the electronic structure of chromophore, which perturbs the electrostatic and hydrophobic interaction within the binding site. This work highlights the critical factors hidden in the protein networks, linking with their experimental photocycle kinetics. It presents an opportunity to quantitatively examine the alternation in chromophore’s equilibrium geometry and identify details which have substantial implications in designing synthetic LOV constructs with desirable photocycle efficiency.

## Introduction

Optogenetics represents a new paradigm that can control biological responses through light. In the past decade, optogenetic tools have rapidly evolved through understanding of the fundamental principles governing the biomolecular systems and technological advancement. These non-invasive tools are promising for diagnosis and treating several diseases afflicting humankind. The advanced application of the optogenetics-based biomedical tool include treatment of diseases such as diabetes, neurological conditions, heart diseases, and cancer.[1] They have also been used for gene-editing and epigenome regulation that have significant applications in molecular and cellular biology.[2] The area of optogenetics could target either neuronal cells that function in neuronal signalling activation or suppression; or non-neuronal cells that affect the signalling cascade for precise monitoring and control of biological functioning. One such example of non-neuronal optogenetic tool development utilizes families of photoactivable proteins - the LOV (light, oxygen, and voltage) family of proteins. LOV domain proteins provide a novel platform owing to their small molecular size and quick response time.[3] These tools take advantage of the fluorescent properties of LOV proteins to develop fluorescence imaging tools and possible implementation in super-resolution microscopy.[4] The LOV-based fluorescent reporter systems have shown increased brightness, improved stability and functionality in low-oxygen and anaerobic conditions, compared to the green fluorescent proteins.[5] In all cases, the core LOV domain signals the effector elements through the highly variable C-ter and N-ter extensions.[6]

The LOV domain is the photosensory module that responds to UVA or blue light[7] and are widespread among several species, including bacteria, fungi, archaea, protists, and plant species. They are a subclass member of the superfamily – PAS domain (Period-ARNT-Singleminded), which comprises α-helical segment and five-stranded antiparallel β-sheets.[8,9] Structurally, the active site of LOV contains a flavin chromophore (Flavin mononucleotide, Flavin adenine dinucleotide, or Riboflavin) and is characterised by the presence of a conserved motif GXNCRFLQ.[8]

The photochemistry of the LOV domain is triggered when the chromophore absorbs light and attains a singlet excited state (S_1_), followed by intersystem crossing to a triplet state (T_1_). The process induces a proton transfer to the N5 atom, changing the hybridization of the C4a atom to sp^3^, and generates a tilt in the planarity of the isoalloxazine ring. This photocycle mediated by a radical-pair mechanism involve the thiol group of a conserved cysteine and the C4a atom of flavin, enabling electron transfer and subsequent formation of a covalent adduct. The ground state (also referred to as the resting or dark state) is characterized by flavin in its oxidized form, absorbing blue light at 450 nm (2.75 eV). The light-adapted signalling state containing the C4a adduct absorbs at wavelength 390 nm (3.18 eV). While the kinetics of the transition, S_1_ → T_1_, occurs in the nanosecond time regime, the conversion of radical pair to adduct falls within microseconds timescale. Notably, the overall photocycle attains thermal reversibility in the dark state,[10] the timescale of this process is a scientific puzzle to be solved as it ranges between a few seconds to several hours in LOV variants.[11] It also raises opportunities to tune the limetime of the photocycle of LOV domain and extend its scope for developing an efficient optogenetic tool.[12] Studies on LOV proteins and their variants have shown that mutation in the chromophore binding site can affect the photocycle duration. In *Avena sativa* phototropin 1 (AsLOV2), mutations like Val to Thr reduces the time constant to a few seconds, Asn to Leu extends to hours, whereas, Asn to Ala has no effect.[13–15] Likewise, in fungal circadian clock photoreceptor Vivid (VVD), Ile to Val double mutants, Met to Ile double mutant, and Met to Leu double mutants reduces, increases, or have no effect on the time decay constant, respectively.[16] The relationship between mutation and changes in the time constant links to the change in electrostatic environment,[17] hydrogen bonding residues[18] and steric effect[19] on flavin affecting the adduct’s stability.[15] Mutations in the active site can perturb the solvent accessibility near the chromophore and facilitate proton transfer involving the photocycle time constants.[20,21]

Several experimental studies based on LOV domain protein structures shed light on the effect of specific amino-acid point mutation, their side-chain orientation, and the potential of being H-bond donors /acceptors in the close vicinity of the chromophore. However, an existing gap in the current pool of knowledge is a comparative structural analysis that could explain the molecular origin of the mutational effect on the LOV photocycle. Here, we have considered several LOV domain proteins from diverse sources to construct a phylogenetic tree and analyzed their clustering pattern. Structural models of protein based on the cluster’s distance are energy minimized and suitable cavities are hydrated with water molecules for structural analysis. Subsequent analyses are based on normal mode simulations to create community networks based on the amino acid arrangement. We also employed graph-based algorithms to understand the H-bonding pattern in LOV proteins and the chromophore binding site arrangement and function.

## Materials and Methods

### Phylogenetic analysis

Given the significant number of LOV protein sequences available in the databases, our foremost approach was to construct a distance-based phylogenetic tree. The objective is to determine their closest relatives among the eukaryotic and prokaryotic origins. However, the phylogeny relationship is based on the LOV domain’s changes in the amino acid sequences. The protein sequences were searched using the NCBI server’s protein Basic Local Alignment and Search Tool.[22] The initial seed proteins were collected from the literature.[23–26], and the search was performed for non-redundant sequences using a threshold E-value of 0.05. The pattern of amino acid and conserved Cys, which forms an adduct with flavin in the light state, is cross-checked for all sequences.

Multiple sequence alignment was performed using the MUSCLE algorithm (See supporting information).[27] Construction of distance trees uses aligned sequences. The MEGA11 program was used to construct the phylogenetic tree employing the maximum likelihood method.[28] Jones-Taylor-Thornton’s substitution model was adopted for amino acids, whereas, the heuristic nearest-neighbour interchange method was used for building the tree inference. To test the phylogeny, we employed the Boostrap method with 1000 replicas. The unrooted distance tree was finally constructed and visualized using the interactive Tree of Life (iTOL) server.[29]

### Structural Modelling

The constructed phylogenetic tree retrieved representative three-dimensional structures from individual clusters from the protein data bank (PDB). In total, 18 such structures constitute the dataset for comparative analysis. The corresponding photocycle kinetics were collected from reported literature. The PDB format of each structure was cleaned to remove any solvent or ion other than protein, a chromophore (FAD, FMN, or Riboflavin), and water molecules. In the case of dimeric and multimeric structures, only one chain is processed for further analysis. Missing residues in the structures are refined using the SWISS-MODEL server.[30] The structural coordinate in 5DJT contains a mutation in the active site Cys50Ala, which was back-mutated to retain Cys in the present work.[31] Note that the resulting network becomes crowded while considering protein residues outside the LOV domain. The resulting network in such a case indicates several nodes that either have one edge or no edge within the path connected to the chromophore. Hence, we only considered the residues present in the PAS domain of LOV to refine the protein network.

The presence of water molecules within the LOV chromophore binding site is crucial for its photochemistry.[20,32] It has been reported to stabilize the internal charges in protein and participate in hydrogen bonding interaction. To account for the hydration of proteins, we used the Dowser++ program for hydrating the biomacromolecule.[33] The prediction of water sites at the cavity is based on an internal probe of 1 Å for finding the cavity and a semi-empirical estimate of binding energy more significant than 10 kcal mol^-1^.

### Hybrid QM/MM calculation

The three-dimensional coordinates of protein structures are input for the hybrid quantum mechanics/molecular mechanics (QM/MM) calculations. The QM region comprises flavin’s isoalloxazine ring and is treated with the TD-DFT method using the B3LYP/6-31G* level of theory. The remaining system constitutes the MM part and is treated with Amber force fields ff14SB.[34] Dispersion correction is applied using Grimme’s D3/B-J damping variant.[35] Other parameters include resolution of identity (RI) for Coulomb integrals (J) and COSX for exchange integrals, denoted as RIJCOSX.[36] In these simulations, the backbone atoms of protein are fixed to their crystallographic position, while side-chain atoms and water molecules present around 8 Å of the isoalloxazine ring are relaxed. All QM/MM-based geometry optimization is performed with Orca program[37] (quantum chemistry package) interfaced with the DL_POLY module (for MM) of Chemshell.[38] The energetics of the structural models are detailed in the supporting information (Table S1).

### Network Construction

Protein networks are constructed based on the QM/MM optimized geometries. The comparative analysis was based on three different network types, viz., (i) hydrogen-bond interaction network, (ii) hydrophobic interaction network; and (iii) residue interaction-based dynamic correlation network.

The residual interaction data obtained from the protein’s normal mode analysis (NMA) displays the residue-by-residue cross-correlations matrix. The preliminary graph consists of amino acids as nodes and the distance between them as edges. It is followed by a coarse-grained approach that uses the Girvan-Newman clustering method. The final network thus converts the all-residue network into communities of highly intra-connected but loosely interconnected nodes.

For simplification of networks, they are represented using centrality measures like degree centrality and betweenness centrality to quantify the relevance of each node in the graph. ‘Degree centrality’ denotes the number of edge incidents to a given node. ‘Betweenness centrality’ refers to the shortest path between two nodes, so the number of edges that pass through is minimized. Notably, the nodes with high betweenness may have considerable influence within a network as they control the information passing between other nodes. Removal of such nodes will disrupt communications as they lie on the most significant number of paths taken by messages transmitted through the network. Interested readers may refer to the study by Deb et al., where similar methods are used to study residue interaction and protein contact networks of LOV proteins.[39] The networks in this study are created using Bio3D working through RStudio IDE, R version 4.2.0.[40] Customization and visualization of networks are performed using the igraph program. Description of protocol for network construction and the codes are shared in the Supporting Information.

The details of hydrogen bonding and hydrophobic interaction are further collected in the form of graph-based networks. The participating residues/water molecules represent the nodes, and the edges link. The connections between nodes are based on Dijkstra’s algorithm. Bridge program, a plugin with Pymol, was used for this analysis.[41] Specific interaction of amino acid with chromophore atoms is analysed using LiBiSCo.[42]

## Results and Discussion

### Sequence-guided phylogenetic reconstruction of LOV proteins

Understanding phylogenetic connections are crucial for a wide range of biological investigations. Major evolutionary transitions rely on an accurate phylogenetic tree, such as molecular adaptation detection, morphological character evolution, and recreating demographic shifts in separated species. Given a large number of LOV protein sequences available in the annotated form, it is crucial to establish a phylogenetic tree based on variation in the protein sequences. Multiple sequence alignment was performed using the Muscle algorithm. The bootstrapped tree was obtained with 1000 replicates as an evolutionary model. The constructed unrooted phylogenetic tree is shown in Figure 1, where the nodes and branches of proteins with similar sequences are clustered together and coloured distinctly. LOV belonging to the same kingdom are often found to have diverse functioning. For instance, LOV protein belong to the same kingdom – Viridiplantae has been classified in to three separate clusters. to the first cluster belong to the plant phototropins, the second cluster contains LOV proteins from zeitlupe (ZTL) and flavin-binding kelch repeat F box protein (FKF1) and the third cluster includes neochromes, aureochromes, and protein twin-LOV.

**Figure 1:**
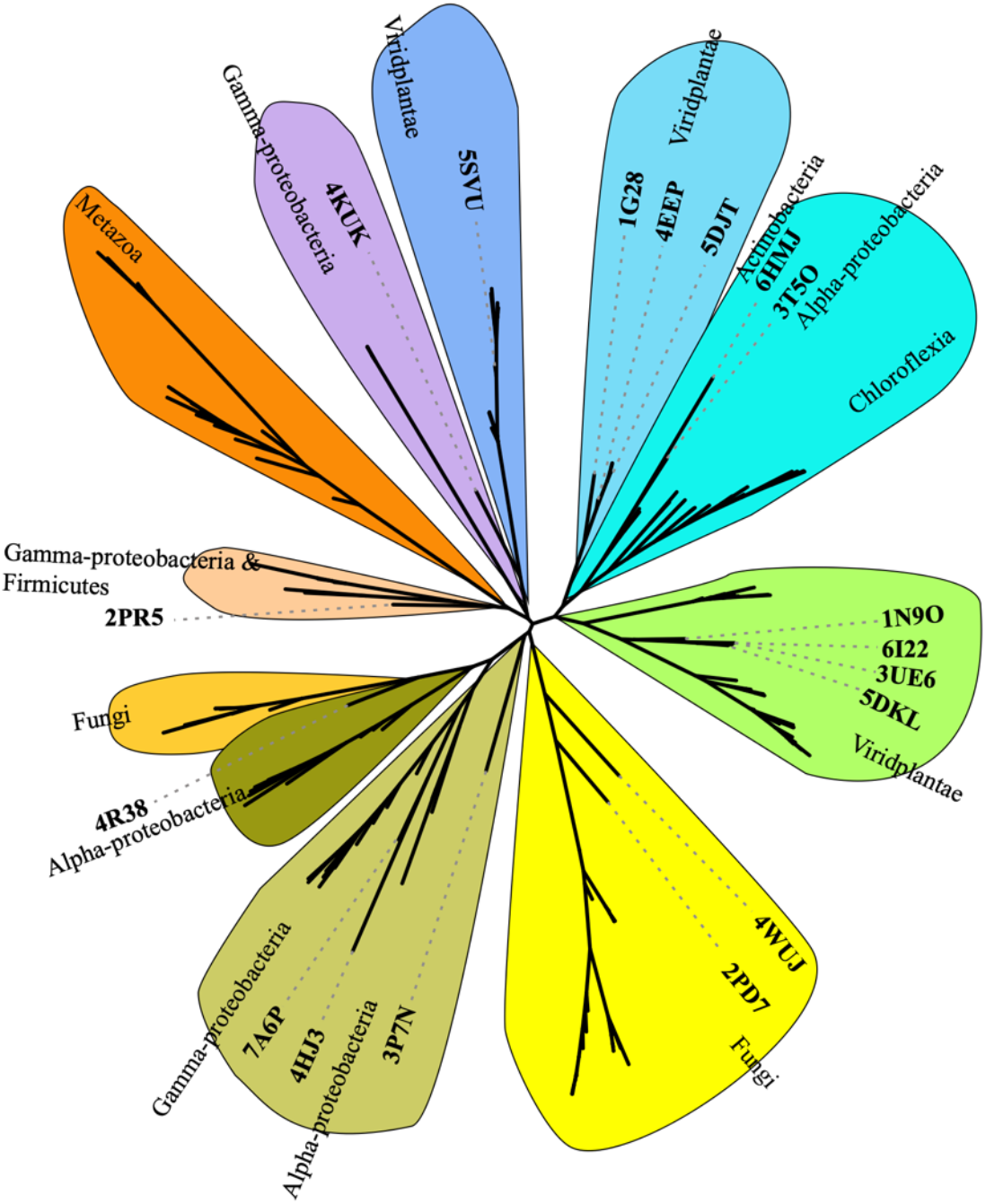
Phylogenetic tree depicting the diversification of the LOV domain proteins.

Another such example is the fungi, which got distributed across two clusters. One class of protein belongs to fungal G-protein receptors, while the other belongs to white collar 1 (WC1) and VVD photoreceptors. Notably, the LOV module is widely dispersed among the proteobacteria belonging to alpha, gamma, and actinobacteria classes. The differences in the sequence and its arrangement in the chromophore binding site are responsible for variability in photoinduced molecular reactivity. Thus, understanding the structural positioning of the amino acid side chains will provide insights on the factors governing the on/off kinetics. Available structural models of the experimentally resolved protein were retrieved from the Protein Data Bank. In total, 18 structural models were considered for network analysis. The corresponding cluster group of these LOV proteins was marked in the phylogenetic tree (Figure 1). More details on the organismal source, along with their rate kinetics and experimental conditions, are provided in the Table 1. For further analysis, the proteins have been categorized into fast, intermediate, and slow kinetics based on their time constants.

**Table 1:** Classification of the proteins based on their time constant values.

| PDB code | Organism | Time constant (and relevant details) | Ref |
| --- | --- | --- | --- |
| 1N9O<sup>a</sup> | *Chlamydomonas reinhardtii* | 200/168 sec | [43] |
| 1G28<sup>a</sup> | *Adiantum capillus-veneris* | 90 sec | [44] |
| 5DJT<sup>a</sup> | *Avena sativa*;<br>*Staphylococcus aureus* | ~50 sec | [17] |
| 4EEP<sup>a</sup> | *Arabidopsis thaliana* | 54/68 sec<sup>d</sup> | [45] |
| 4KUK<sup>a</sup> | *Dinoroseobacter shibae* | 9.6 sec (20 °C) | [46] |
| 7A6P<sup>a</sup> | *Pseudomonas putida* | 210 sec (20 °C) | [47] |
| 3UE6<sup>a</sup> | *Vaucheria frigida* | 480 sec | [48] |
| 3P7N<sup>a</sup> | *Erythrobacter litoralis* | 25.5 sec (25 °C, pH 6) | [49] |
| 4HJ3<sup>b</sup> | *Cereibacter sphaeroides* | ~39 min | [50] |
| 2PR5<sup>b</sup> | *Bacillus subtilis* | ~65 min (23 °C, pH 7) | [51] |
| 6HMJ<sup>b</sup> | *Nakamurella multipartita* | ~37 min (22 °C) | [52] |
| 4R38<sup>b</sup> | *Erythrobacter litoralis* | ~55 min | [53] |
| 4WUJ<sup>b</sup> | *Trichoderma reesei* | ~25 min | [54] |
| 5DKL<sup>b</sup> | *Phaeodactylum tricornutum* | ~25 min | [55] |
| 6I22<sup>b</sup> | *Ochromonas danica* | 112 min<sup>d</sup> | [56] |
| 5SVU<sup>c</sup> | *Arabidopsis thaliana* | ~6.6 hrs | [57] |
| 3T50<sup>c</sup> | *Brucella melitensis* | ~9.5 hrs (20 °C, 6.5 pH)<sup>d</sup><br>~5.8 hrs (35 °C, 7 pH)<sup>d</sup> | [58] |
| 2PD7<sup>c</sup> | *Neurospora crassa* | ~2.7 hrs (25 °C)<sup>d</sup> | [59] |

**Notes:** <sup>a</sup>fast kinetics (<500 sec); <sup>b</sup> intermediate kinetics (500 to 7000 sec; up to 112 min); <sup>c</sup> slow kinetics (>7000 sec; >2 hrs); <sup>d</sup> referred to as half-life in original reports.

### Classification of data set based on the time constant

The structural models constituting the dataset in this study are classified based on the adduct decay (time constant) kinetics. In particular, three regimes of photocycle kinetics are used for classification, viz. fast (<500 sec), intermediate (500 to 7000 sec), and slow (>7000 sec) (detailed in Table 1). The protein networks constructed in our work are based on normal mode analysis, aiming to understand the chain connectivity related to the folding pattern of proteins. Processing the coordinate data from PDB with coarse-graining helps decompose constituent amino acid residues’ mutual and intrinsic motion. Here the centrality of individual residues represents the topological importance of the residue-residue cross-correlation network. The residues act as nodes and the edge which interconnect two nodes are weighted in proportion to the degree of correlated motion. The betweenness centrality in the network was calculated by finding the shortest path between pairs and scoring each edge (n) of the paths with the inverse value of the number of shortest paths (m); betweenness score = n/m. Note that the residues present at the C-ter and N-ter of model proteins did not show a direct perturbative effect on the chromophore binding site but indirectly through allosteric pathways. Since the total number of residues in each protein structure is not constant, it is challenging to identify a particular trend from the resulted protein network. For structural modelling, the terminal residues were removed to bring the protein networks within a range of betweenness centrality. Thus, the shortest paths for all pairs were optimized according to the classification of the dataset. ranging between 14 to 17 for fast cycling, 20 to 24 for intermediate cycling, and 30 to 33 for slow cycling proteins. Details of individual protein models are provided in Table 2. A graphical representation of the classification based on betweenness centrality is shown in Figure 2. It also includes the total number of residues in each protein of the structural dataset. The coordinates of modelled structures of the LOV domain used for constructing networks are available in the supporting information.

**Table 2:** Details of LOV structural models used for network construction.

| Protein Name | PDB code | N-ter residue<sup>a</sup> | C-ter residue<sup>a</sup> | Total residue number<sup>b</sup> | Water molecules added<sup>c</sup> | Centrality<sup>d</sup> |
| --- | --- | --- | --- | --- | --- | --- |
| Phot-LOV1 domain | 1N9O | 4His | 109Ser | 106 | 25 | 17 |
| Flavin-binding domain, LOV2, from the chimeric phytochrome/phototropin photoreceptor PHY3 | 1G28 | 1Lys | 104Met | 104 | 9 | 17 |
| LOV2 Domain of Avena Sativa | 5DJT | 13Asn | 121Asp | 109 | 15 | 16 |
| LOV2 domain of phototropin 2 | 4EEP | 6Ile | 115Val | 110 | 23 | 15 |
| A superfast recovering full-length LOV protein from the marine phototrophic bacterium | 4KUK | 20Asp | 139Leu | 120 | 17 | 16 |
| Short LOV protein PpSB2-LOV | 7A6P | 31Val | 146Glu | 116 | 16 | 16 |
| Blue-light photoreceptor Aureochrome1 LOV | 3UE6 | 46Gln | 154Ala | 109 | 12 | 14 |
| Light-activated transcription factor El222 | 3P7N<sup>e</sup> | 72Ser | 177Glu | 106 | 22 | 12 |
| LOV protein | 4HJ3 | 14Asp | 127Ser | 114 | 15 | 23 |
| Dimeric LOV Photosensor YtvA | 2PR5 | 10Val | 117Tyr | 108 | 25 | 20 |
| RNA-binding Light-Oxygen-Voltage Receptor | 6HMJ | 243His | 350Val | 108 | 19 | 21 |
| EL346 blue-light-activated histidine kinase | 4R38 | 28Ile | 129Gln | 102 | 9 | 20 |
| Fungal LOV Domain Photoreceptor | 4WUJ | 37Ser | 146Glu | 110 | 27 | 24 |
| Blue light photoreceptor Aureochrome 1a LOV | 5DKL | 17Gln | 128Val | 112 | 10 | 20 |
| LOV domain | 6I22 | 19Val | 123Tyr | 105 | 19 | 20 |
| LOV domain of ZEITLUPE | 5SVU | 14Thr | 133Ile | 120 | 24 | 33 |
| LOV-HK sensory protein | 3T50 | 2Ala | 116Val | 115 | 15 | 33 |
| Fungal Blue-Light Photoreceptor Vivid | 2PD7 | 27Val | 148Cys | 122 | 19 | 30 |

**Notes:** <sup>a</sup> Residue numbers corresponds to the reference sequence in ‘Protein Feature View’ of PDB. <sup>b</sup> Total number of amino acids present in the structural model used for normal mode analysis. <sup>c</sup> Number of water molecules added in the 8 Å of isoalloxazine ring, using the Dowser program. <sup>d</sup> Betweenness Centrality.

**Figure 2:**
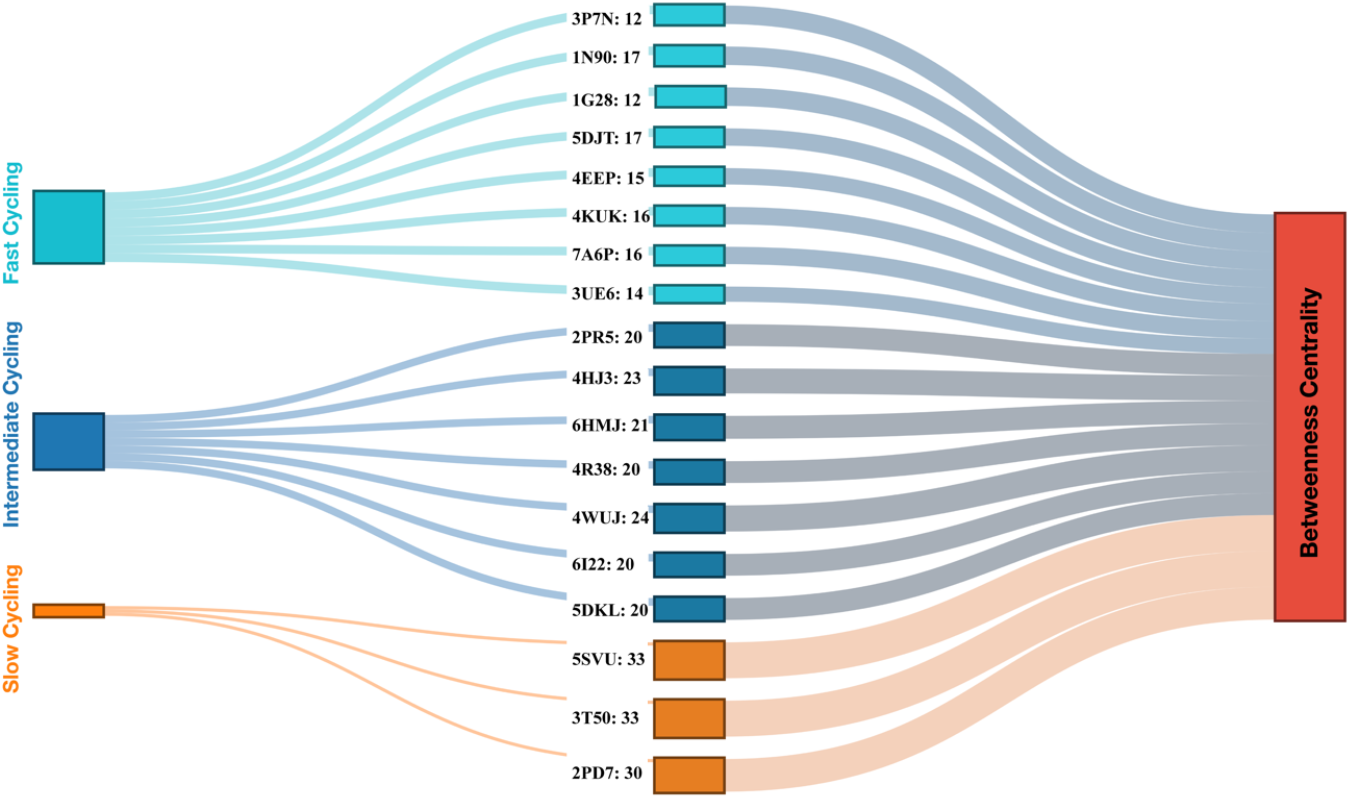
Dataset classification LOV domain based on adduct decay kinetics and betweenness centrality of protein networks.

### Interaction analysis of the chromophore within the active site

Interaction details help quantify the factors responsible for the stabilization of chromophore. Figure 3 shows the interaction details of LOV proteins exhibiting fast kinetics. The ring I in the isoalloxazine was involved in polar contacts with the electronegative N and O atoms. On the contrary, ring III engages in hydrophobic interaction, attributed to the presence of steric bulky methyl groups. The presence of H-bonding of the C2=O group in the active site accelerates the photocycle kinetics.[60] It also stabilizes the isoalloxazine ring within the active site, as revealed through vibrational frequency measurements.[61] Except for 7A6P, 5DJT, and 4KUK, the rest of the fast kinetic class of proteins showed water molecules to be in close contact with the carbonyl moiety adding to its polarity.[62] (Figure 3F, G, and H). Within the active site, the polar contact is also facilitated through Asn, as observed for all fast kinetics proteins. In 4EEP the Asn residue is making polar contact with C4=O, which however could re-orient during the dynamic conformational changes (Figure 3E). The dynamic flipping of Leu and Gln residues stabilizes isoalloxazine from both sides.[63] Interestingly, another Asn residue is involved in making H-bond with N3 and/or C4=O group. In the case of IG28, 3UE6, 4EEP, and 5DJT, we observe an interaction between a Gln residue and the N5 of ring II (Figure 3A, B, E, and G). Previous reports have identified this residue crucial for tuning the LOV photochemistry inferred from mutation studies.[59,64,65] In the absence of Gln residue in 3P7N, 1N9O, and 7A6P, the H-bonding within the active site are compensated by water molecule(s) (Figure 3C, D, and F). The contour map of hydrophilic interaction also shows substantial contributions from Arg and other Gln residues. However, their interactions are not associated with the isoalloxazine ring but with the phosphate chain and are crucial for chromophore stabilization. Arg residue forms a conserved salt bridge with the phosphate group and reduces the dynamicity of chromophore.[63] Note that this interaction has also been predicted to stabilize adducts, by affecting the decay kinetics of the adduct.[66] Overall, we see the interactions of Gln/Asn residues and water molecules are predominantly electrostatic in nature and interacts with ring I as they contains polar atoms. Whereas, the hydrophobic Val, Leu, and Ile residues interacted adjacent to ring III containing two methyl groups. The Ala residues found in a few structural models, are known to facilitate faster kinetics.[62] Some of the aromatic-aromatic contacts (π-π stacking type) mediated through Phe and Tyr residues are also observed.

**Figure 3:**
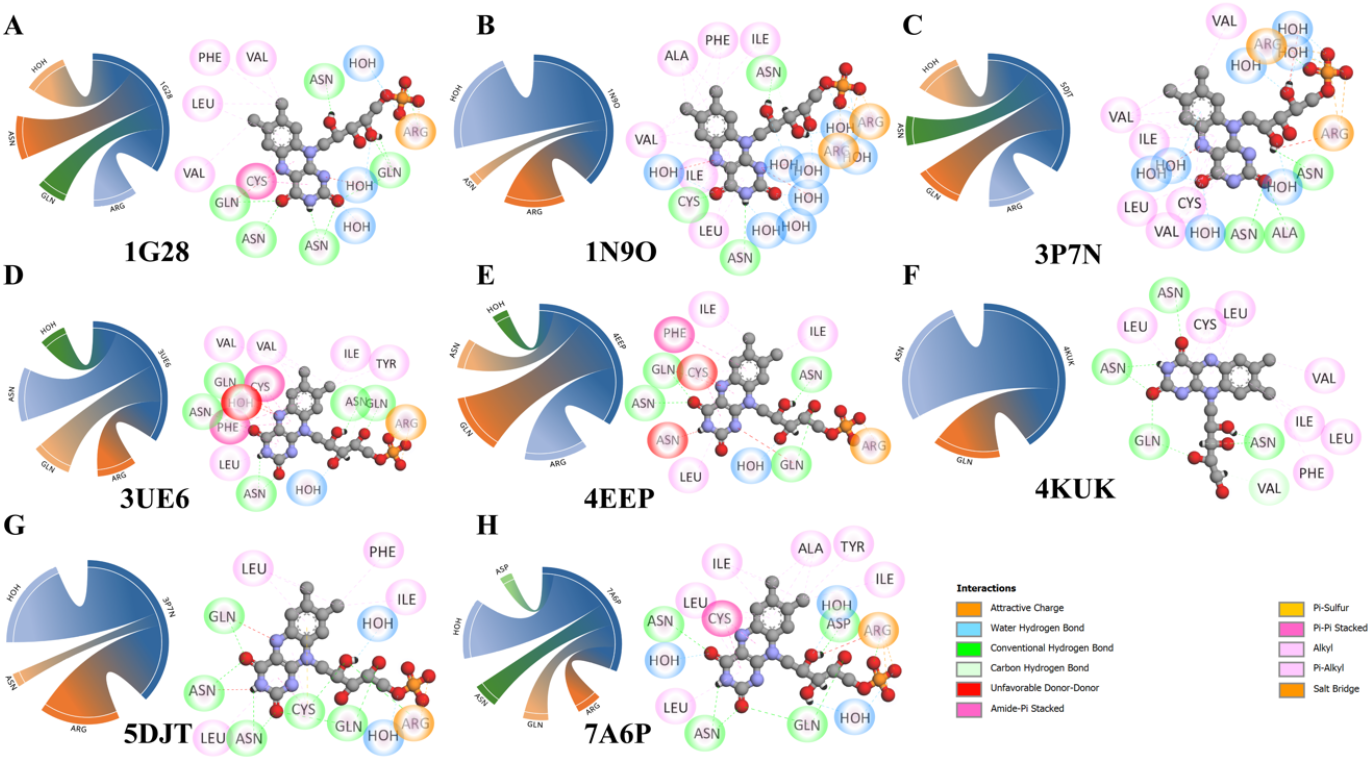
Interaction analysis of fast-cycling LOV proteins (A-H). The contour map shows the relative weight of polar contacts (polar interaction). Residue-wise interaction is shown for each protein model with chromophore in stick model and residues coloured according to the nature of hydrophobic and polar contacts. The colour codes used to describe interaction are shown.

In the case of intermediate cycling proteins (Figure 4A-G), one Asn residue was involved in making H-bonding with C2=O. Asn residue stabilizes the chromophore interaction by forming polar contact with N3 and/or C4=O. Exceptions to this trend are 6I22 and 4WUJ (Figure 4D and E). The role of Gln residue in stabilizing the isoalloxazine ring through polar contacts is evident except for 6HMJ (Figure 4G). The role of water molecules in relaying the H-bond to the isoalloxazine ring at N5 position is a well-established hypothesis.[67–69] A water molecule close to N5 is in line to this for the intermediate cycling proteins, except 4R38 and 4WUJ (Figure 4C and E). Notably, the extent of polar contacts of the ring-I is relatively less than that of the fast kinetic group. Additional water-mediated contacts strongly involve hydroxy groups in the extended phosphate chain. Arg residue forms salt-bridges with phosphate and Gln and/or Asn residues are involved with H-bond with the hydroxy groups, rendering the chromophore more stable. The hydrophobic contacts involving the π-alkyl and π-π stacking type interactions are similar to that of the fast kinetic group. Interestingly, the Cys and Met in this group prefer π-Sulfur type interaction, instead of induced-dipole interactions.

**Figure 4:**
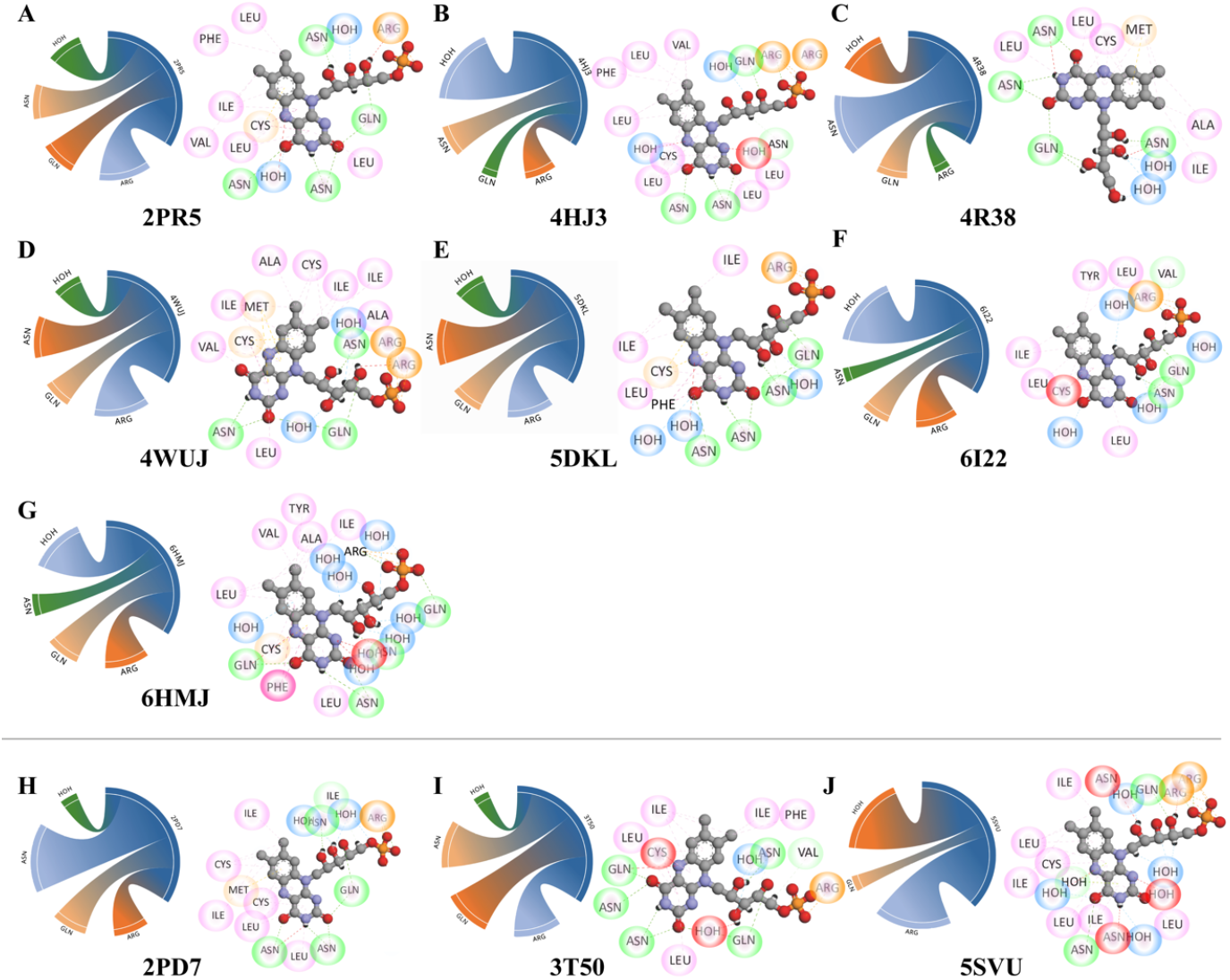
Interaction analysis of intermediate cycling (A-G) and slow cycling (H-J) LOV proteins. The contour map shows the relative weight of polar contacts (polar interaction). Residue-wise interaction is shown for each protein model with chromophore in stick model and residues coloured according to the nature of hydrophobic and polar contacts. The colour codes used to describe interaction are the same as in Figure 3.

The number of structural models in the slow kinetic groups is comparatively less, making it difficult to identify a reliable trend. Nevertheless, the water molecules mediating stabilization through H-bond with hydroxyl groups of extended chains exists only in 2PD7 and 3T50 (Figure 4H and I). In 5SVU (Figure 4J), one additional water molecule enabled H-bond with N3 of ring I and N5 of ring II. The role of Asn in forming conventional H-bonds is consistently observed in this group, and Gln is found only in the case of 3T50. The hydrophobic interactions were predominantly mediated by Leu and Ile residues through π-alkyl type contacts with ring III. The hydrophobic interaction involving a larger side chain or conversely, the absence of shorter side chain like Val, could be a contributing factor for longer time scale in decay.[62]

### Quantifying interacting with chromophore atoms

This section quantitatively illustrates the residue-specific interaction with atoms of the chromophore. Figure 5 shows the average distance between active-site residues with specific atoms of the isoalloxazine ring of flavin. The H-bonds are mainly formed by three polar atoms, in which nitrogen at N3 and carbonyl groups’ oxygen at O2 and O4 contribute significantly. Considering the three kinetic groups, the average distance between N3, O2, and O4 with the corresponding residue follows a similar trend. The distance is in order O4 > O2 > O4 (Figure 5A, C, and D). However, the contributions with water molecules are not include included in this analysis. We also performed distance analysis for hydrophobic interactions analysis. The pattern of average interaction distance obtained shows differences crucial for their photocycle kinetics. The N5 atom found the minimum distance in all three groups, indicating that strong hydrophobic interaction is incident at the chromophore’s reaction centre. For the fast kinetic group (Figure 5B), relatively distant interaction is for C6 and C9a. On the contrary, the maximum interaction distance for the intermediate kinetic group is for C9a and C9, whereas for slow, it is C9 and C7M (Figure 5D and F). This indicates that the dominant factor determining the photocycle kinetics is the hydrophobic and steric restraint induced due to non-polar residues in the active site.

**Figure 5:**
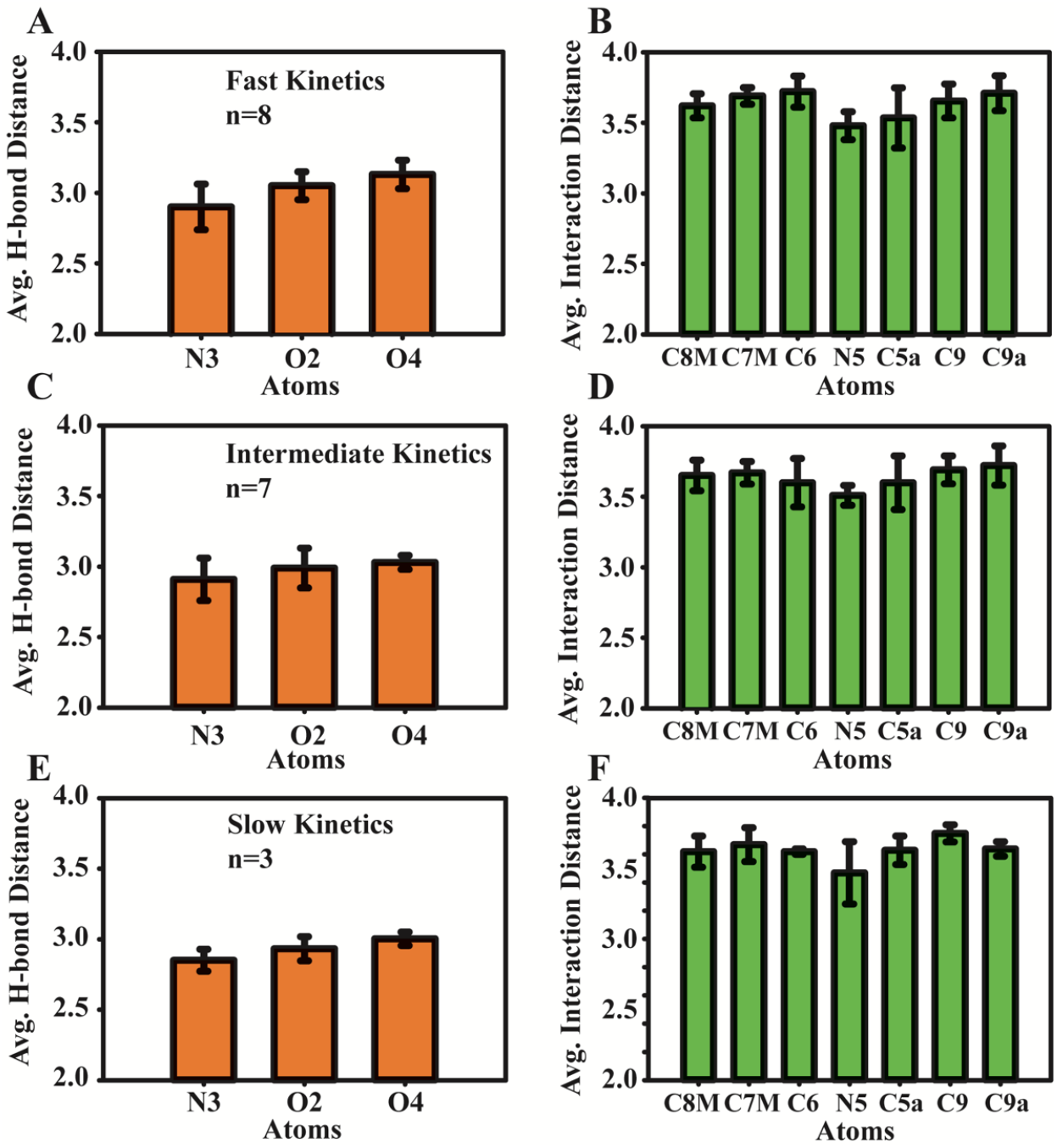
Average interaction distance between chromophore atoms and active site residues are shown for fast (A, B); intermediate (C, D); and slow kinetic group (E, F). H-bond distance on left panel and interaction distance on right panel denotes polar and hydrophobic interaction, respectively.

### Dynamic correlation network

The dynamic correlation network was constructed using the information from normal mode analysis. The residue-based network was converted into community clusters and weighted according to their significance within the network. Calculation of these undirected network graphs considers the betweenness centrality for partitioning them into clusters.[40] The networks are optimized for a range of betweenness centrality, corresponding to their classification according to kinetics. It helps in avoiding unambiguity and rationalizes the networks for comparison. Tight packing of chromophore within the LOV binding site is facilitated by β-sheets surrounded by Eα and Fα helices. Their cross-talk between community have direct effects on the stability of the adduct. It is evidenced through a network of H-bonds and electrostatic interaction, facilitated mainly by residues present in Eα, Hβ and hydrophobic by Gβ. While residues in Iβ contribute through both polar and non-polar contacts.[9,46] The protein correlation network for the fast cycling group are in Figure 6, where the community clusters are shown with distinct colours. The corresponding protein segments are rendered with similar colours to denote the network nodes.

**Figure 6:**
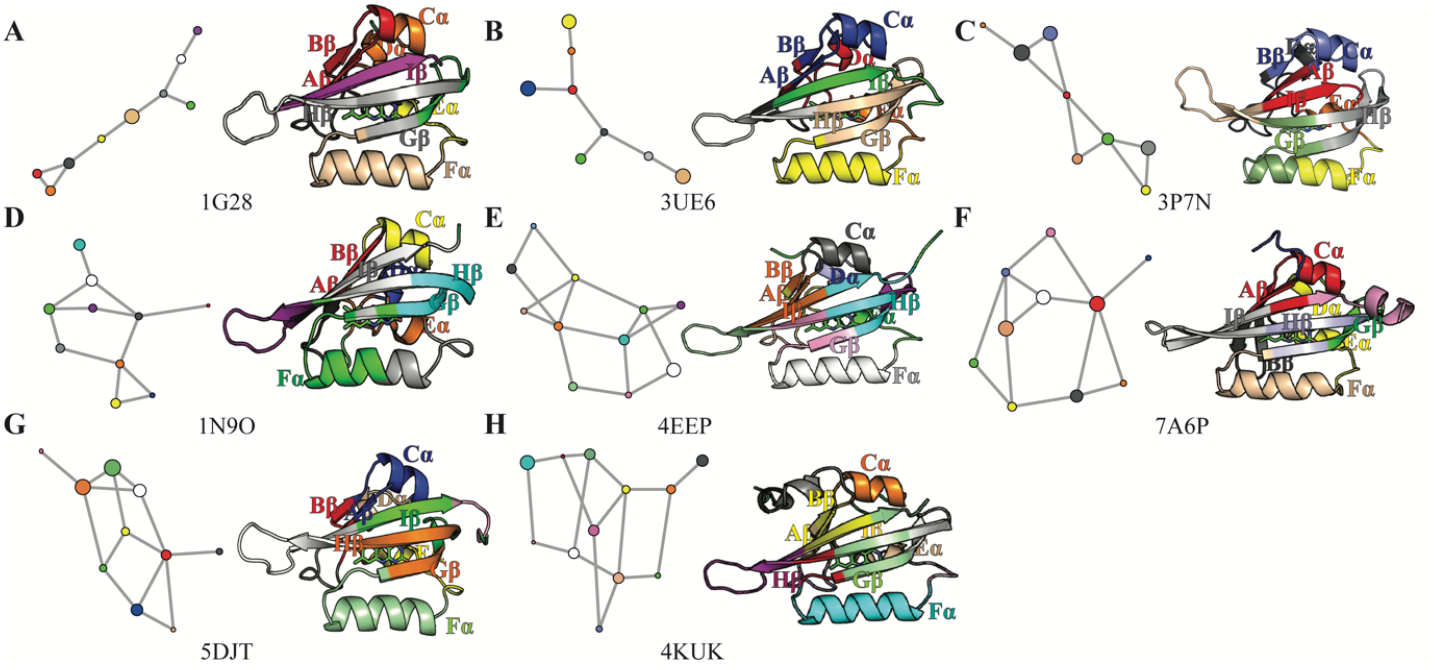
Dynamic correlation network of fast-cycling LOV proteins (A-H). The spheres represent coarse-grained networks based on dynamically coupled communities, and the linking is based on calculated betweenness. Similar colours represent individual community clusters on the correlation network and protein’s secondary structures.

The graph pattern from 1G28 and 3UE6 was devoid of interlinked branches, except for two short bifurcations (Figure 6A, B). The number of nodes present in these networks are 9 and 8, respectively. The β-sheets (Gβ and Hβ) are intensely engaged in community interaction inall structural models of this group, with an exception of 3UE6. Likewise, Iβ also show strong network interaction except for 3UE6 and 1G28. The loop regions between β-sheets, connecting Bβ-Cα, and Eα-Fα were found to be central and are linked to two or more nodes in the dynamic correlation network. Conformational changes in these regions alter photocycle kinetics as (i) they are solvent-exposed on one side with buried hydrophobic part; (ii) they are reported to show temperature dependency.[8,44] The correlation network of 1N9O and 7A6P are alike, containing 10 nodes (Figure 6D, F). The phototropin 1N9O and bacterial protein 7A6P can accommodate more water molecules within the chromophore binding site.[44,47] The triad in the network which links Fα with the loop between Hβ-Iβ and connecting the β-sheets can facilitate increased solvent accessibility. This promote faster deprotonation (flavin N5 position) and faster kinetics of 200 and 210 sec (Table 1). The number of nodes present in the correlation network of 3P7N are 8, whereas, 5DJT contain 10 nodes. Notably, the number of nodes are high in 4KUK and 4EEP containing 12 and 11 nodes, respectively.

The hydrophobic region in β-sheets is crucial for LOV proteins having intermediate kinetics. In the case of 2PR5, Hβ and Iβ denote the central node connected to several residual communities (Figure 7A).[70] The congregation formed in 4R38 through β-sheets and Fα appears dispersed but are connected to the more residual community. They are crucial in governing collective normal modes. Though the time constants for 2PR5 and 4R38 are comparable (3880 and 3300 sec), the key difference could be the presence of chromophore and riboflavin, respectively. Moreover, the faster adduct decay in 2PR5 (Figure 7C) are due to factors like - the presence of Ala in Fα instead of conserved basic residues lacks a polar contact with an extended chain; the presence of residues Met in re-face; and Phe near isoalloxazine ring.[51,53] The Gβ-Hβ-Iβ region is linking Aβ-Bβ in one closed loop and to Fα in another loop, within the network of 4HJ3 (Figure 7B), restricting the solvent accessibility towards chromophore site. Ile and Leu residues, positioned re-faced to isoalloxazine ring, govern the solvent perturbation and the kinetics.[50] The total number of nodes in the network of 2PR5 and 4HJ3 are 11, while it is 9 in 4R38. The Iβ sheet in 4WUJ loosely packs the chromophore site, indicating the possibility of solvent exposure towards the active site (Figure 7D). This hypothesis is based on collected evidence from force-field-based simulations.[69] The corresponding residual community is linked to Aβ and Bβ, and is collectively crucial for governing the decay kinetics.[24] The network pattern found in 5DKL and 6I22 are alike, where the part of Fα with Gβ, Hβ, and Iβ appear as central nodes (Figure 7E and G), playing a crucial role in chromophore stability. The simulation-based machine-learning analysis confirmed the dynamic difference between these regions and the protein’s remaining part.[71] Notably, the hydrophobic residues in Iβ region are essential for cross-talk between other residual communities of protein.[72,73] The number of nodes revealed in the dynamic correlation network for 4WUJ, 5DKL, 6HMJ, and 6I22 are 10. Notably, the involvement of Aβ, Bβ, and Cα remains uniform in all protein models. They are known to participate in solvent mediated polar contact on the outside and facilitating a hydrophobic environment towards (inside to) the chromophore. However, the role of Fα in cross-talk with β-sheets is more significant in the intermediate kinetics group.

**Figure 7:**
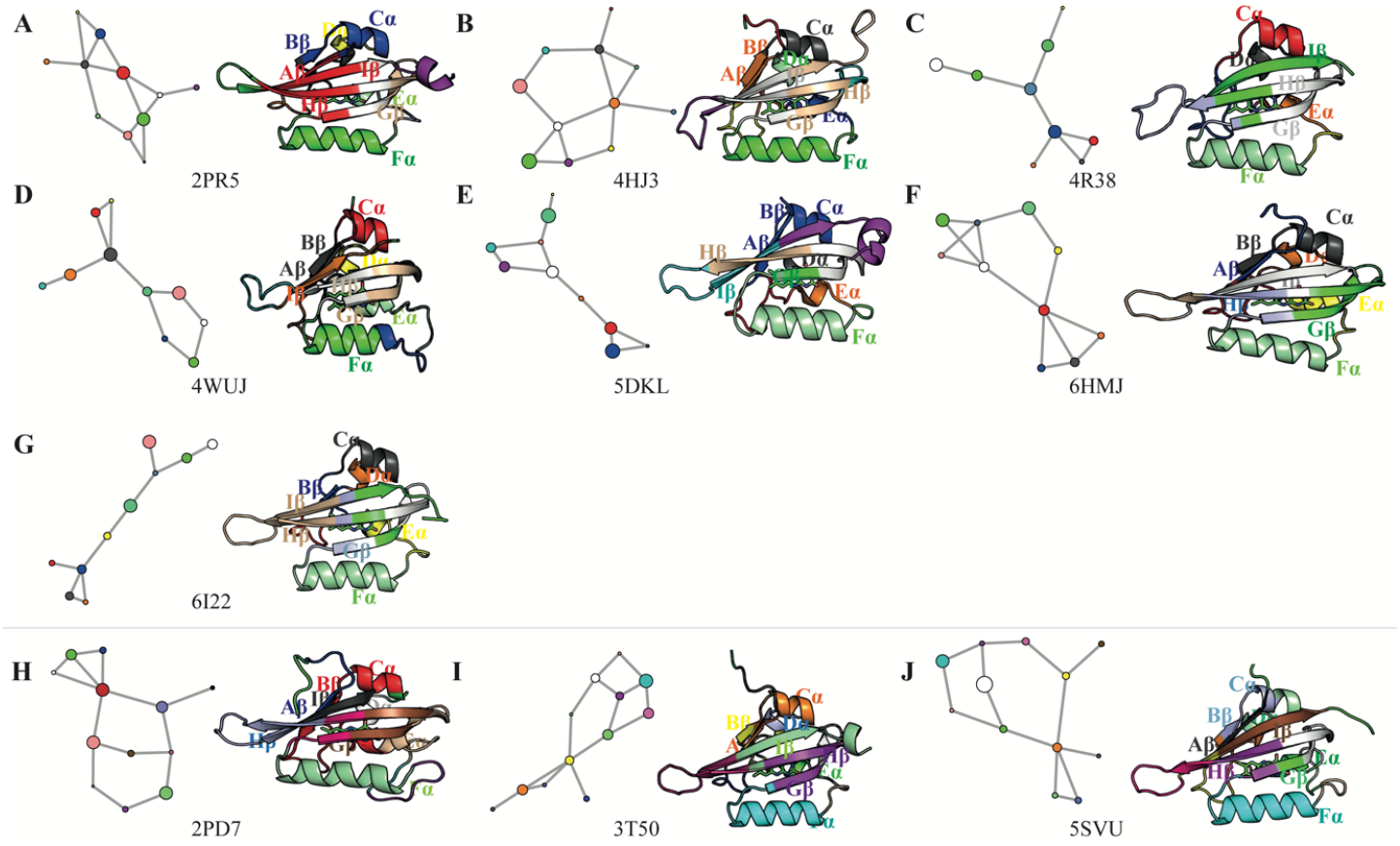
Dynamic correlation network of intermediate (A-G) and fast (H-J) cycling LOV proteins. The spheres represent a coarse-grained network based on dynamically coupled communities, and the linking is based on calculated betweenness. Similar colours describe individual community clusters on correlation networks and protein’s secondary structures.

All three protein models of slow kinetics, viz. 2PD7, 3T50 and 5SVU have 12 nodes in the complex dynamic correlation network. The role of Bβ is crucial in connecting with other residual communities and maintaining close packing of the chromophore in all three models. In the case of 2PD7, perturbation in Bβ-loop-Cα is critical in maintaining the polar contacts by Gln and Asp (Figure 7H).[59,74] Met and Arg role in stabilizing the sheet-helix packing has also been demonstrated through molecular dynamics simulation.[75] Notably, in 3T50, the sheet-loop-helix region at N-ter remains central to other adjacent loops of the residual community (Figure 7I). The structural compactness of these regions remains the same after adduct formation, as determined through NMR chemical shifts.[58] However, in the case of 5SVU, this region is shared by more than one node (Figure 7J). Overall, the Bβ-loop-Cα region appears crucial for slow kinetic LOV protein; but more structural models are required to understand its significance.

### Correlation between bond length and rate kinetics

The Hybrid QM/MM method helps evaluate the protein environment’s effect on the chromophore, especially on the bonded and non-bonded interactions.[76] Particularly for flavin chromophore, this method has proven instrumental for geometry optimization and calculation of spectroscopic properties.[77] The structural models of the LOV domain are segmented into three groups based on their rate kinetics and subjected to hybrid QM/MM simulation to identify the changes in equilibrium bond length of chromophore. The QM region selection was uniform in all models, which constitutes the isoalloxazine ring. The protein environment was treated as point charges for which the parameters were obtained from the MM force fields. We analyzed the standard deviation in bond lengths over the structural models for each bond. The numerical data obtained from simulations are summarized in Table S1 of Supplementary information, and the graphical representation is shown in Figure 8. Note that the residues proximal to the chromophore directly impact its geometry. The corresponding sequence logos for each dataset are shown in Figure 8A by comparing active site sequences for all proteins in Figure 8B-D.

**Figure 8:**
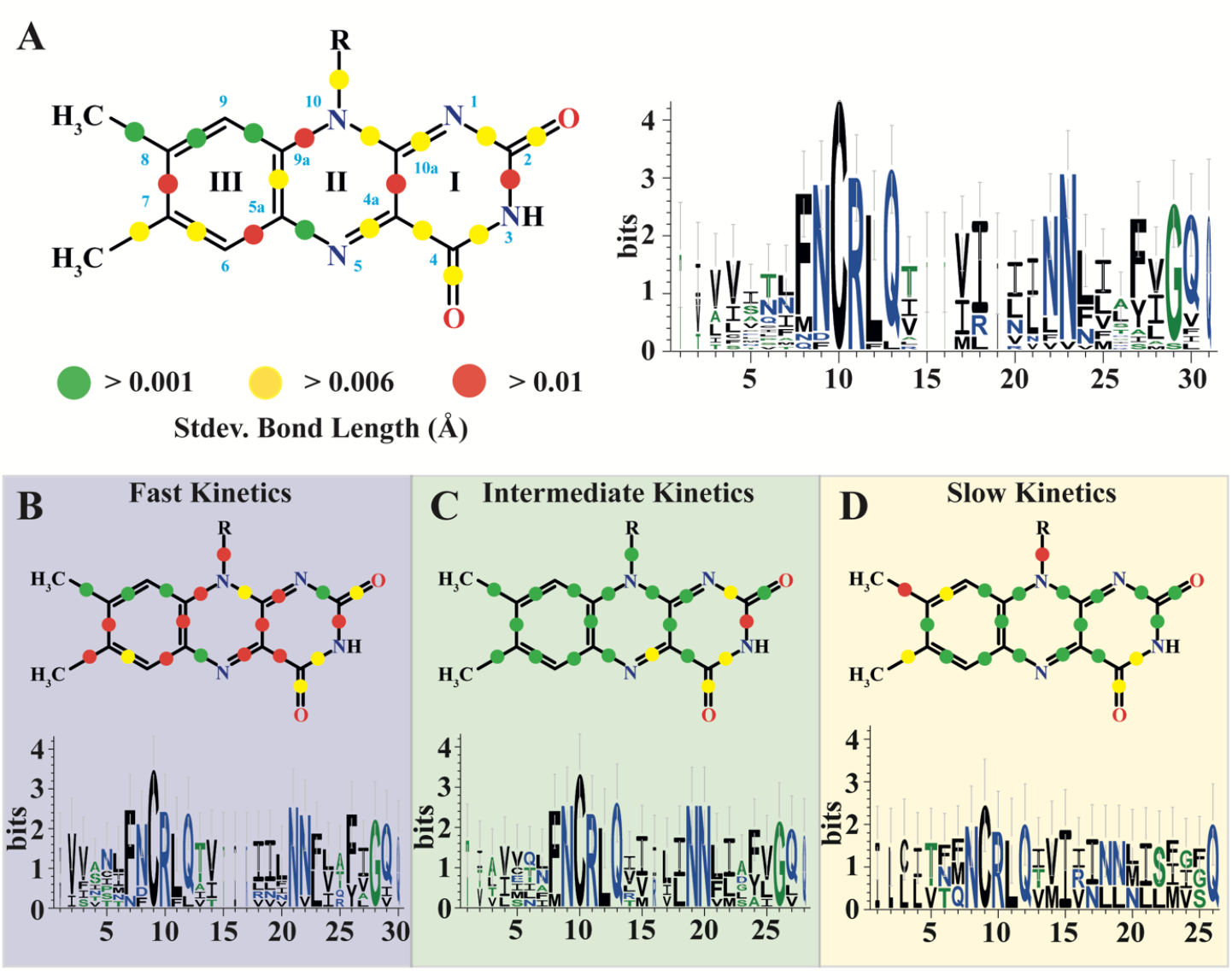
Alternation in the equilibrium geometry of chromophore due to rearrangement of polar and hydrophobic interaction within the active site. The standard deviation in bond lengths was obtained based on hybrid QM/MM calculation. Colour codes used to denote alternation include green (>0.001 Å), yellow (>0.006 Å), and red (>0.01 Å). The sequence logo represents the multiple sequence alignment of active site residues within 6 Å of the chromophore. The height of individual residue at a particular position corresponds to its relative frequency. The figure panels correspond to all structural models (A), fast kinetic (B), average kinetic (C), and slow kinetics (D) groups.

The collective analysis of protein models (Figure 8A) depicts that the least affected bond lengths (>0.001 Å) are CH_3_-C8, C8-C9, C9-C9a, and C5a-N5, which are linked to dimethylbenzimidazole moiety (ring III) of the isoalloxazine ring. On the contrary, the regions where bond length vary significantly (>0.01 Å) are prevalent across the three rings. Two bonds, viz. C2-N3 and C4a-C10a are part of ring I, and the remaining three C5a-C6, C7-C8, and C9a-N10, belonging to hydrophobic groups are distributed across ring III. Among the bonds constituting an electronegative element (N or O atom) in the rings I and II show intermediary bond length deviation (>0.006 Å) with C4-C4a being an exception. However, this pattern differed significantly when the individual groups were analysed based on their experimental rate kinetics. Substantial differences were found in the bond length of the isoalloxazine ring for the three kinetics groups.

The kinetics and the protein environment at the binding site correlated well with the changes in the bond lengths of the chromophore (Figure 8B-D). Significant structural changes were observed for the chromophores belonging to fast kinetics. Compared to the collective analysis, the deviation in bond lengths is now altered from intermediate to high deviation (std. dev.), as reflected by the change in contour colour from yellow to red (Figure 8A and B). The bonds concerning the diazabutadiene system (N1=C10a-C4a=N5) and fused rings II and III (C9a=C5a). The highest perturbation of ring I, in which two more bonds, viz. N1-C2 and C4-C4a show alteration from intermediate to least (green) and high (red), respectively. Overall, the isoalloxazine ring shows substantial alterations, implicating that steric constraints by the active site residues are less onto the chromophore binding site in this group. On the contrary, the remaining two groups (Figure 8C and D) reflect the minimum perturbation on the chromophore. The bonds linked with a ring I show deviation in the group of proteins showing intermediate kinetics. The bond involving polar elements offers intermediate changes, which includes N1-C2, N3-C4, C4=O, and C4a-N5, and remains consistent with cumulative analysis. Likewise, C2-N3 shows significant alteration (>0.01 Å), same as that of the fast kinetics group. For the slow kinetics group, the isoalloxazine ring has the most negligible perturbation in bond length as shown by the cumulative analysis. Exceptions include C8-CH_3_ and C8-C9 of ring III, which show high and intermediate deviation. Overall, the bonds involving electron-rich N5 are altered >0.01 Å in the case of fast, >0.006 Å in intermediate, and >0.001 Å in slow kinetics.

### Sequence logo analysis with specific chromophore atoms

We extended our analysis for chromophore’s atom-wise interaction with active site residues (Figure 9). The occurrence of amino acids interacting with particular atoms of the chromophore is shown as a sequence logo, where the height of the letter is proportional to its frequency. The average distance between N3, O2, and O4 for H-bond interactions (ring I) and C8methyl, C7methyl, C6, C5a, C9, and C9a (ring III) are shown in Figure 5. The list also includes hydrophobic interaction mediates through N5 (ring III). Particularly for H-bonding, Asn and Glu are key players, which is also reflected in Figure 8. However, their differential impact has been clearly understood from the QM/MM simulations.

**Figure 9:**
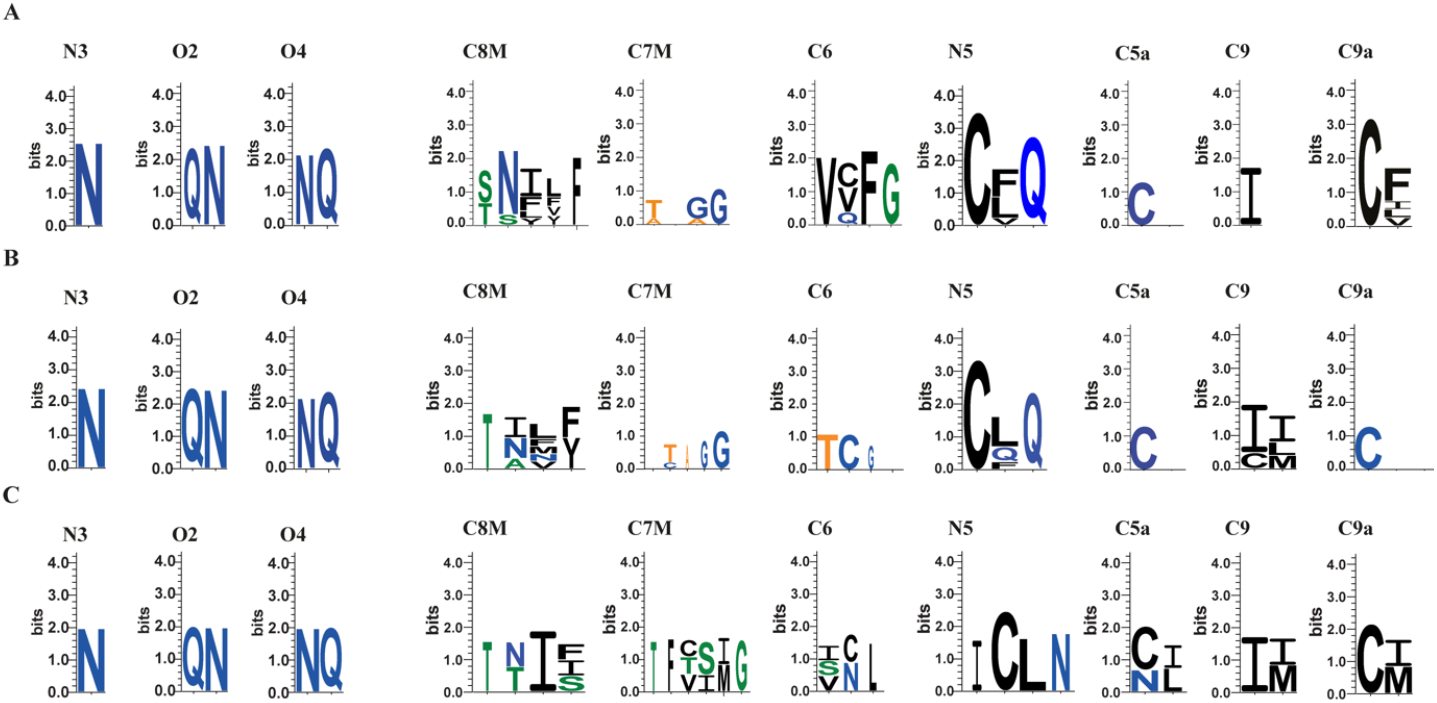
Sequence logo analysis for amino acids interacting with individual atoms of chromophore. The analysis from three kinetics group are shown for (A) fast, (B) intermediate, and (C) slow. N3, O2, and O4 analysis represent the H-bonding interaction while the remaining atoms describes the hydrophobic interaction.

The C8M region of isoalloxazine’s ring III is highly occupied with several amino acids. The interaction occupancy is nearly similar in fast and intermediate, whereas it is strongly dominated by Ile and is occupied with fewer amino acids in slow kinetics. On the contrary, the interacting amino acids near C7M are significant for slow kinetics proteins compared to the other two groups. For the C6 atom, strong hydrophobic restraints are present in proteins with fast kinetics and are minimum for the intermediate. The adjacent atom C5a shows the involvement of Cys residue, which, however, is occupied with other residues only in the case of the slow group. The hydrophobic interaction near the reaction centre N5 was found to be equivalent for the fast and intermediate groups, with a slight difference in the frequency of amino acids. On the other side (near the ribityl chain), the C9 atom interacts only with Ile in the fast group. At the same time, positional restraints are reflective in the other two groups with different amino acids, such as Cys, Leu, Ile, and Met. The C9a atom is also occupied with Cys interaction, especially in the intermediate group. Notably, this atom is occupied with another hydrophobic residue in the fast and slow groups. Thus, the variable hydrophobic interactions near the chromophore are critical for alternating equilibrium bond length. Later they translate it into photocycle kinetics, which provides crucial information for designing synthetic LOV constructs.

## Conclusion

The translational utility of the LOV domain as sensors is limited due to insufficient information on the role of the local protein environment on the photocycle kinetics. Mutational studies reveals that polar group for mediating electrostatic interaction, hydrophobic side chain residues in the proximity of chromophore, and water molecule accessibility, are the factors that perturb the LOV photocycle kinetics. On top of focusing on individual residues, analysis of residual clusters around chromophore also provides a strong lead for constructing synthetic proteins. In this work, we revisited the local protein environment using similar approaches. However, we limited our structural analysis to the dark-state conformational models of LOV protein. One reason is their wide occurrence in monomeric conformation in the dark state. In contrast, the light-activate state in certain classes of proteins attains a dimeric conformation and operated in more complex protein networks.

Our analysis emphasises the role of hydrogen bonding near ring I and the hydrophobic environment surrounding ring III, which cooperates with the chemical nature of the isoalloxazine ring. The role of Asn, Leu, Gln, and water molecules are rendered common among the LOV protein having fast photocycle kinetics. For other kinetic groups, the role of Val, Leu, and Ile in governing hydrophobic interaction and Arg in mediating salt-bridge with phosphate backbone remains common. Notably, the absence of shorter sidechain residue is responsible for a longer decay time constant. Dynamic correlation network analysis reveals that the α-helices and β-sheets links are critical for restricting water accessibility near N5 of Flavin. Cross-talk between the Fα with the Hβ-Iβ loop and adjacent β-sheets is also crucial in restricting water accessibility near N5 of Flavin and affect the kinetics. The role of Aβ, Bβ, and Cα are non-significant in fast and intermediate proteins. Tight packing of the Bβ-Cα region is responsible for slow kinetics. Based on quantum mechanics, we found a significant difference in Flavin’s equilibrium geometry that correlates well with the experimental time constant. Taken together, future studies comprising the larger structural dataset can render a more accurate correlation between protein networks and photocycle kinetics.

## CRediT authorship contribution statement

**Rishab Panda:** Conceptualization, Investigation, Writing – original draft, Software **Pritam K. Panda:** Software, Interpretation. **Janarthanan Krishnamoorthy:** Reviewing and Editing, **Rajiv K. Kar:** Supervision, Investigation, Writing-original draft, Reviewing and Editing.

## Acknowledgement

This research was supported by Department of Science and Technology (DST) – Science and Engineering Research Board (SERB), Government of India (SRG/2022/000858) to RKK. The author also thanks ICMR Centre for Excellence, Grant no. 5/3/8/20/2019-ITR; INUP, MeitY (5(1)/2021-NANO); and RnDOPs, IITG for a research grant (Project reference no. 2022-040113000301).

